# Persistence of remnant boreal plants in the Chiricahua Mountains, southern Arizona

**DOI:** 10.1101/2020.03.02.974055

**Authors:** Anda Fescenko, James A. Downer, Ilja Fescenko

## Abstract

Boreal plants growing along southern edge of their range on isolated mountains in a hot desert matrix live near the extreme of their physiological tolerance. Such plants are considered to be sensitive to small changes in climate. We coupled field observations (1974, 1993, 2019) about the abundance and vigor of small populations of ten remnant boreal plant species persisting in uppermost elevation spruce-fir forests of the Chiricahua Mountains, together with a theoretical modeling of the species’ tolerances to three climate change cues: warming, drought, and forest fire, in order to explore the persistence of frontier boreal plant species in the frame of climate changes. We hypothesized that populations of these cryophilic plants have declined or become locally extinct during an adverse warming period since 1993, enforced by two large forest fires (1994, 2011). We used plant functional traits and principal component analysis to model tolerances of the plants to combined actions of warming, drought, and forest fire. Our model predicted selective sensitivity to warming for two species: *Vaccinium myrtillus* and *Rubus parviflorus*, while possible decline of the other species could be explained by drought and/or fire. We surveyed the study area in 2019 and found eight of the ten species still occur in the area. Five species occurred in wet canyons at lower elevations, but three species persisted in low vigor at the uppermost elevation highly affected by fires. Both warming-sensitive species did not show signs of decline: population of *R. parviflorus* increased in abundance and vigor, while *V. myrtillus* persists without significant changes since 1993. Despite the recorded increase in temperature in the study area over one degree Celsius between years 1975-1993 and 1994-2019, our study did not find evidence of the direct warming effect on the observed species. We conclude that severe wildfires and multi-decadal decrease in precipitation rather than warming are the main limiting factors of the remnant boreal species’ remarkable but limited persistence in the Chiricahua Mountains. Our study demonstrates how field observations can be combined with modeling to evaluate species selective responses to different environmental stresses for better environmental management decisions, particularly in light of climate change.

## Introduction

During the last few decades, researchers and society are increasingly worrying about climate warming impact on species diversity and survival (1–4). Some remnant boreal plants growing along the southern edge of their range and/or at the uppermost elevations are expected to be heralds of adverse climate change (5, 6). Plants growing on isolated mountains in a hot desert matrix are often near the extreme of their physiological tolerance and may be sensitive to small gradual changes in temperature (7). Warming in mountains is accompanied not only by extreme heat waves, drought, and severe forest fires (8), but also by invasion pressure from lower-elevation plant communities (9). However, plant species evolving under continuous climate fluctuations and persistent stresses, have evolved strategies allowing them to survive such environmental changes. Such strategies include temporal persistence in refugium – for rapid climate changes (10), shifting elevation or geographical range (migration) – for slower changes (9, 11, 12), and physiological adaptations – for slow climate changes if elevation or geographical range are limited (13, 14). Surveying of remnant boreal plants growing near the extreme of their physiological tolerance could help both estimate pressure of climate change on the biosphere and promote better understanding and protection of potential refuge areas for endangered species.

In European mountains, where plant survey traditions started more than 400 years ago (15), plant communities at the uppermost elevations are well studied. Many studies have been carried out on the impact of recent climate change on plant abundance and diversity, e.g., (16, 17). These studies brought some expected and unexpected results, such as: elevation upshifting and downshifting for different plant groups (16–19); increase of local species richness (16, 20, 21); and decrease of rare or endemic boreal plants (16, 22). In the North American Southwest, boreal plants persist on the summits of isolated mountains – ‘sky islands’ – within a hot desert matrix that significantly limits their possibilities for adjusting to elevation and geographical range. Such desert islands are under-recognized natural laboratories for testing plant-climate interactions. As far back as in 1974, a set of remnant boreal plants was surveyed in the Chiricahua Mountains, southeastern Arizona (23). The abundance and vigor of ten perennial boreal species persistent in the spruce-fir forests were recorded along five hiking trails around Chiricahua Peak. The second survey was done 19 years later (24), when most populations of surveyed species were found at 1974 levels or had increased in number. After a large forest fire in 1994, most of the populations were suspected gone, see postscript in (24).

In this study, we performed the third field survey (2019) of the same boreal plant species. The aim of our study was to test a hypothesis that populations of these cryophilic plants have declined or become locally extinct during an adverse warming period since 1993 and following extensive fire disturbances. We surveyed the abundance and vigor of the ten plant species along the same hiking trails around Chiricahua Peak. Fire severity was estimated from field observations, while multi-decadal changes in annual temperature and precipitation were calculated from local climatic data. We performed univariate and multivariate analyses to model each species tolerances to the three anticipated adverse environmental cues (stress factors) of climatic change: warming, drought, and forest fire. The species were divided into functional groups according to their likelihood to be affected by one stress factor (selective sensitivity). In this paper, we report changes in the remnant species’ abundance and vigor during the period 1994–2019 and explain them by using a functional trait approach within the framework of climate changes. Our study reintroduces the historical surveys performed by Moir in 1974 and 1993 and paves the road for systematic and regular surveys of species that are sensitive to climate change in the North American Southwest.

## Materials and methods

### Study area

The Chiricahua Mountains are located in southeastern Arizona (Figure 1) and are the biggest range of the Madrean Sky Island Archipelago in the United States. The range is approximately 65 km long and 32 km wide, with maximum elevation of 2976 m a.s.l. (Bennett et al. 1996). The upper reaches of the range are dominated by a series of ridges and peaks in excess of 2700 m a.s.l. The local climate is characterized by bimodal precipitation with a considerable range in daily and seasonal air temperature (25). Approximately two-thirds of annual precipitation falls during summer thunderstorms, the remaining precipitation falls in winter as snow (25). Common trees at elevation above 2400 m a.s.l. are Douglas fir (*Pseudotsuga menziesii*), Engelmann spruce (*Picea engelmannii*), southwestern white pine *(Pinus strobiformis)*, quaking aspen (*Populus tremuloides*), white fir (*Abies concolor*), *Arizona pine* (*Pinus ponderosa* var. *arizonica*).

**Fig. 1.**
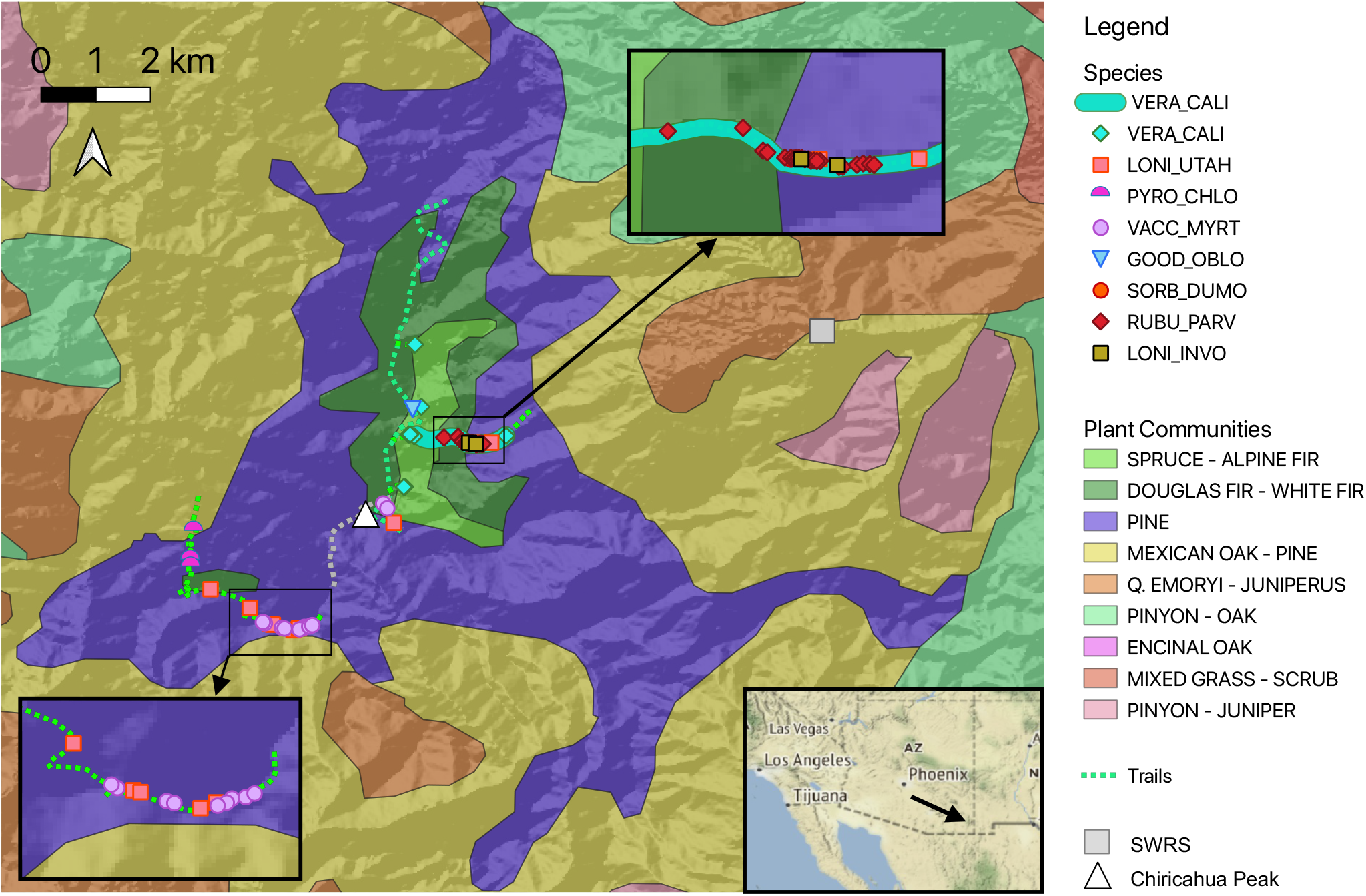
Study area in the Chiricahua Mountains, southeastern Arizona: plant communities (AZGEO) and locations of the observed plant species in 2019. For species codes see Table 1. Surveyed trails shown by green dashed lines. SWRS – Southwestern Research Station, Portal, Arizona. The inset in lower right corner shows location of the Chiricahua Mountain range in Arizona, USA.

The study area is located at elevations of 2400 to 2976 meters, where the spruce-fir forests occur (Figure 1). Note that the large-scale map of plant communities does not show little patches of spruce-fir forests dispersed along the trails. The area was severely affected by two of the largest recorded wildfires in the Chiricahua Mountains: Rattlesnake Fire (11’129 ha forested area burned in June and July of 1994), and Horseshoe 2 Fire (90’226 ha burned in 2011) (26). The fires burned at various intensities across almost all upper elevation forests including all of the study area.

### Studied species and field survey

Ten boreal plant species were studied: *Chimaphila umbellata* (Ericaceae; Prince’s Pine), *Erigeron scopulinus* (Asteraceae; Rock Fleabane), *Goodyera oblongifolia* (Orchidaceae; Western Rattlesnake Plantain), *Lonicera involucrata* (Caprifoliaceae; Black Twinberry), *Lonicera utahensis* (Caprifoliaceae; Red Twinberry), *Pyrola chlorantha* (Ericaceae; Greenflowered Wintergreen), *Rubus parviflorus* (Rosaceae; Thimbleberry), *Sorbus dumosa* (Rosaceae; Arizona Mountain Ash), *Vaccinium myrtillus* (Ericaceae; Bilberry), *Veratrum californicum* (Melanthiaceae; California Corn Lily) (Table 1). These species occur in the uppermost elevation (> 2400 ma. s.l.) of the Chiricahua Mountains and most of them are growing along the southernmost edge of their distributions. The exception is *Chimaphila umbellata*, which subspecies *C. umbellata* subsp. *acuta* has its main distribution area much more southward (in Mexico). This species was surveyed in our study to be consistent with previous surveys. Eight of the ten species are North American plants, while *Vaccinium myrtillus* and *Pyrola chlorantha* are circumboreal. Two species – *Erigeron scopulinus* and *Sorbus dumosa* – are endemic to the southwest of North America.

**Table 1.** Abundance, vigor and habitat fire severity of ten boreal plant species in 2019. Plant abundance recorded for each species by counting the number of plots with a particular species, the number of locations, and the number of individuals/ramets (see for details Methods). Values of species vigor and habitat fire severity per plot were averaged over all plots of each species. For plant vigor and habitat fire severity classes see Methods and Tables S2 and S4

| Species | Species code | Number of |  |  | Vigor | Fire severity |
| --- | --- | --- | --- | --- | --- | --- |
|  |  | plots | locations | individuals |  |  |
| *Chimaphila umbellata* | CHIM_UMBE | 0 | 0 | 0 | - | - |
| *Erigeron scopulinus* | ERIG_SCOP | 0 | 0 | 0 | - | - |
| *Goodyera oblongifolia* | GOOD_OBLO | 1 | 1 | 7 | 1.5 | 3.2 |
| *Lonicera involucrata* | LONI_INVO | 2 | 2 | 6 | 2.8 | 3 |
| *Lonicera utahensis* | LONI_UTAH | 9 | 7 | 18 | 1.6 | 3.4 |
| *Pyrola chlorantha* | PYRO_CHLO | 3 | 3 | 9 | 1.8 | 2.8 |
| *Rubus parviflorus* | RUBU_PARV | 22 | 6 | 259 | 1.8 | 2.8 |
| *Sorbus dumosa* | SORB_DUMO | 1 | 1 | 1 | 1.8 | 2.8 |
| *Vaccinium myrtillus* | VACC_MYRT | 11 | 5 | 90 | 1.8 | 2.8 |
| *Veratrum californicum* | VERA_CALI | 54 | 5 | 389 | 1.8 | 2.8 |

Plant species were inventoried along five hiking trails in September-October 2019. The surveyed trails were: Crest Trail, upper part of the Greenhouse Trail, Monte Vista Lookout Trail, and adjacent trails, where spruce-fir forests occurred (see Figure 1, and Table S1 for description of trails). Plants were observed in a 10-m band on both sides of trails (5 meters to each side). The total observation area was of 20.5 km length and 10 m width. A 10-m-long plot was assigned, when at least one of the surveyed species was encountered. That way the size of each plot was 10 × 10 m. In total, 98 plots were assessed. Observations of abundance (number of individuals or ramets) and vigor for each species in each plot were recorded. Vigor of plant was assessed in three classes by evaluating shoot robustness, flowering/fruiting capacity, and symptoms of disease/herbivory/insects/parasitism: low vigor (1), medium vigor (2), high vigor (3) (see for the vigor classes Table S1). Mean species vigor value from all plots, where individuals of a particular species occur, was taken to evaluate the species vigor. In addition, forest fire severity in the observation plots was estimated in five classes according (11): unburned (1), scorched (2), light (3), moderate or severe surface burn (4), deep burning or crown fire (5) (see for the fire severity classes Table S3). Mean fire severity value of all plots, where individuals of a particular species occur, was taken to evaluate the species habitat fire severity. In order to compare our data with historical data, the term ‘location’ was also introduced and adjacent plots with the same species were counted as one location. Therefore, species abundance data were expressed in three levels: as a number of plots with a particular species, as a number of individuals/ramets, as well as a number of locations (only for comparison with the historical data). Locations of plots were recorded by taking geo-referenced photos with a smartphone (accuracy < 30 m). Specimens of the seven studied species were also collected for the SWRS (Southwestern Research Station) Herbarium (see for the herbarium data Sup. Note 4).

### Data processing and analysis

Daily temperature and precipitation data for the years 1956–2019 (measured at the SWRS weather station, Portal, Arizona) were obtained from the GHCN (Global Historical Climatology Network) server.

The weather data were averaged each year as a time series in Wolfram Language; then multidecadal mean values of three time segments: 1956–1974, 1975–1993, and 1994–2019 (corresponding to the field survey years) were compared. Field data were tested for ecological relevance by calculating Pearson correlation coefficients between observed species abundance, vigor and habitat fire severity. Elevations of the observed plants were obtained in QGIS by matching plant locations points with an elevation map (Aster Global Digital Elevation Map).

Plant functional traits were used to estimate possible impact of the three environmental cues of the climate change – increasing temperature (warming), decreasing precipitation (drought), and extreme climatic events (forest fire) – on the surveyed plants. Species tolerances to fire and to drought were obtained from trait databases, while species tolerance to warming was derived from traits directly related to the species response to temperature extremes. We used a set of four traits related to the lower (Minimum hardness zone, Minimum temperature) and the upper (Maximum hardness zone, Minimum elevation in Arizona) limits of the species’ temperature ranges to estimate the species’ tolerances to warming. We also used two additional traits (Plant moisture-use type, Minimum precipitation) to improve values of the plant tolerance to drought. All traits were obtained from the global plant trait database TRY (27), except elevation data that were obtained from SEINet data portal (28), and hardiness zones stemming from the PFAF database (29). Traits of *Sorbus scopulina* were used if those of *Sorbus dumosa* were undefined. When multiple traits were used, the means of scaled numerical values were taken to estimate tolerances. We used principal component analysis (PCA, dudi.pca in ade4 package, R language) to model species’ tolerances to the combined actions of warming, drought, and fire.

## Results

### Local climate change estimate

The weather station (PORTAL 4 SW) is located at elevation 1644 m a.s.l. less than 6 km northeast from the plots (see Figure 1). There was a significant rise of mean temperature of 1.2°C between years 1975-1993 and 1994-2019 (Figure 2a) that is consistent with 0.27°C per decade reported in (30). Note that our data is not adjusted for the time of observation bias (31). The dataset reports a shift of local observation time made in 1991 from 18:00 to 17:00 – towards larger warming drift (31) that could be responsible for the sudden change of the temperature trend in 1991. The weather data shows, first, an increase (in 1975–1993), and then a significant decrease (in 1994–2019) in precipitation during the observed period (Figure 2b). This decrease falls into the long-term oscillating drought pattern of the Southwest (32). We did not find a significant shift in local snowfall (Figure 2c). There was, however, a significant downward shift in mean snow depth in 1994–2019 (Figure 2d) that coincides with a period of human-caused large wildfires occurred in the area and thus may reflect the impact of the fires. Pearson correlation *ρ* between mean annual snow depth and mean annual precipitation is *ρ* = 0.44, but between mean annual temperature and precipitation is *ρ* = −0.15.

**Fig. 2.**
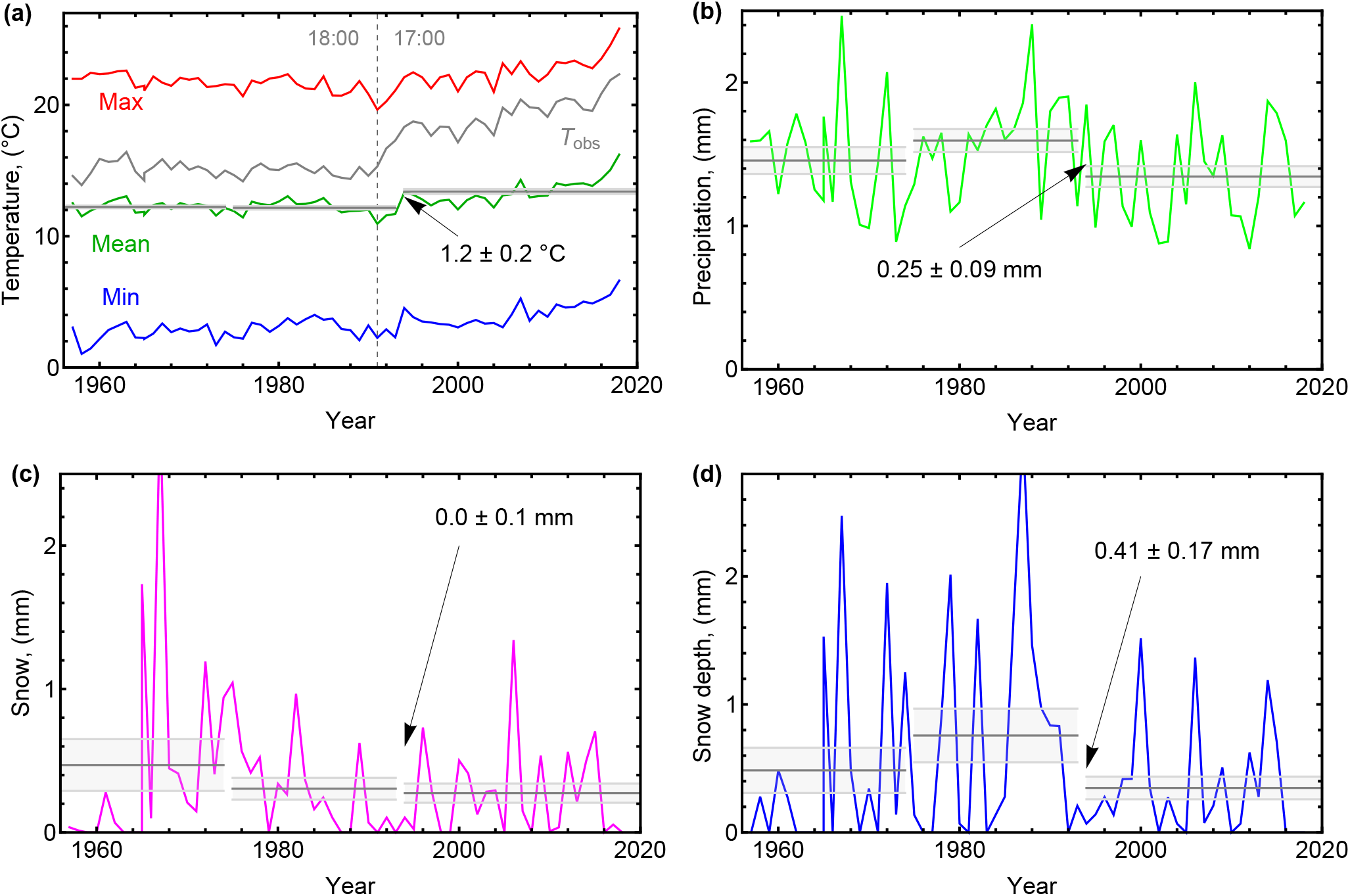
Weather data of the PORTAL 4 SW weather station, averaged per year. Multi-decadal means (for years 1956–1974, 1975–1993, 1994–2019) are shown by gray horizontal lines with standard mean errors intervals (lighter lines). Numbers are values of multi-decadal shifts shown with standard errors. **(a)** Average annual temperatures. *T*_obs_ is the temperature at the time of observation. The vertical dashed line shows the year when the time of the observation was changed. **(b)**Average annual precipitation. **(c)** Average annual snowfall. **(d)** Average annual snow depth. See annual mean values in Table S4.

### Field observations

In 2019, we found eight of the ten focal species (Table 1). In five plots, more than one of surveyed species were found. The most abundant and the most vigorous were populations of *Veratrum californicum* (found in 54 plots, average vigor 2.9) and *Rubus parviflorus* (22 plots, average vigor 2.5). *Vaccinium myrtillus* was relatively abundant, but their shoots were weak and sparse, and some of them had only a few leaves (about 4–6 per shoot) We did not observe any *Chimaphila umbellata* and *Erigeron scopulinus*, and we found only one individual of *Sorbus dumosa*,which was in a poor condition (vigor 1). Species’ habitat fire severity varied from scorched (2) to deep burning/crown fire (5). None of the plots were unaffected by fire. Populations of *Vaccinium myrtillus* were observed most frequently in areas where fire damage was severe (average fire severity 4) and populations of *Veratrum californicum* occurred in places with the least fire impact (average habitat fire severity 2.5). Abundance of the observed species positively correlated with species vigor (Pearson correlation *ρ* = 0.58) and negatively correlated with habitat fire severity (*ρ* = −0.44; Figure 3a). The correlation between species vigor and habitat fire severity was *ρ* = −0.68. *Goodyera oblongifolia* and *Vaccinium myrtillus* occupied the uppermost elevations (higher than 2800 m a.s.l.; Figure 3b); *Pyrola chlorantha* grew at the lowest elevations and at one location even below 2200 m a.s.l., i.e., at the lowest edge were spruce-fir forests occur (see Figure 1). Individuals of *Lonicera utahensis* observed at the lower edge of its elevation range had low vigor. For detailed descriptions, locations, and photos of species see Sup. Note 4.

**Fig. 3.**
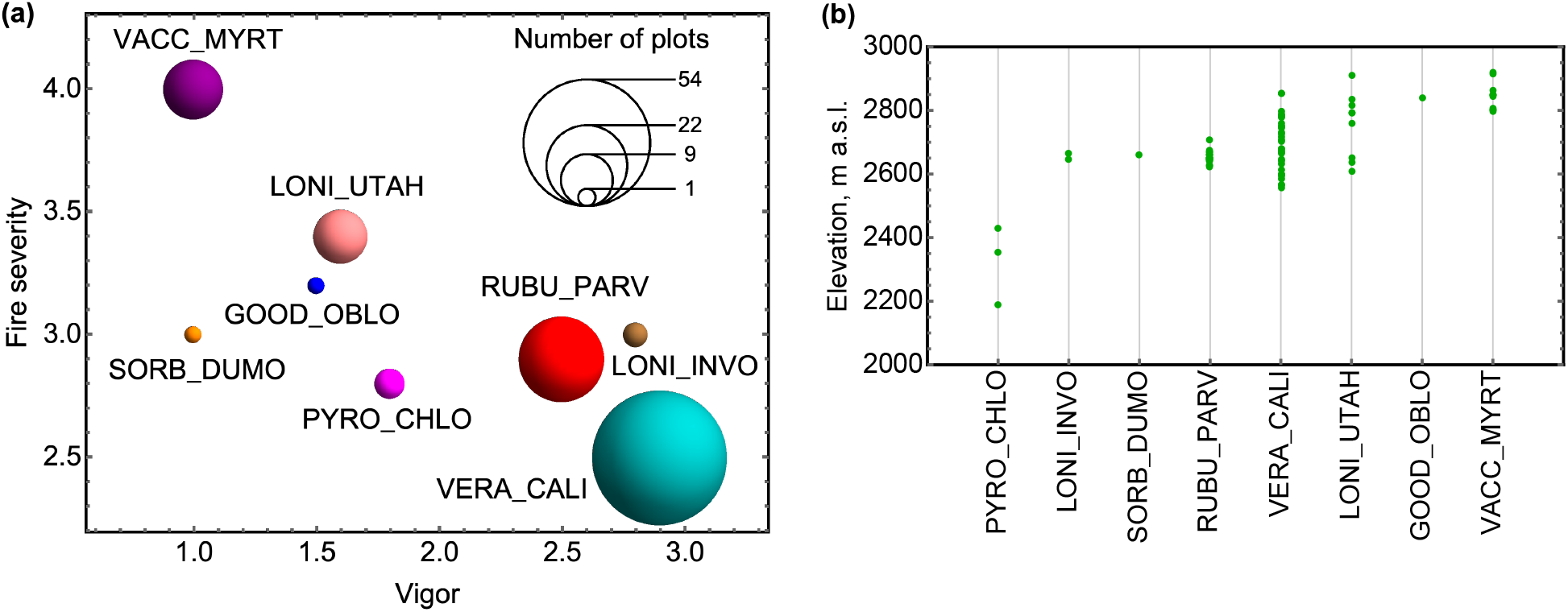
**(a)** Species abundance (number of plots with a particular species) versus habitat fire severity (averaged per species) and species vigor (averaged per species). **(b)** Elevation of the observed species (plots) in meters above sea level. For species codes see Table 1.

### Trait-based model of species’ tolerances to climate change

Since 1993, plants in the Chiricahua Mountains have experienced two large forest fires and gradual decrease of precipitation (see Figure 2b). To check the impact of warming on the plant survival, these two environmental cues (variables) were also considered in our study. Figure 4a shows scaled tolerances of the surveyed plant species to warming, drought, and fire, estimated as described in Methods. Figure 4b shows PCA of the species negative response (intolerance) to the combined effect of all the three environmental cues. Based on the PCA results the plants are divided into four functional groups. The first group includes *Erigeron scopulinus, Pyrola chlorantha*, and *Chimaphila umbellata* that are less affected by any of the studied environmental cues. However, the division for *Erigeron scopulinus* might be rough since functional trait data for this species were limited. The second group includes *Sorbus dumosa* and *Goodyera oblongifolia* that are affected primarily by the large forest fires. The third group includes both *Lonicera* species and *Veratrum californicum* that are more sensitive to precipitation decrease (and fire) than to the warming. Finally, the fourth group includes *Vaccinium myrtillus* and *Rubus parviflorus* that are of special interest in this study since these species are more sensitive to warming but less sensitive to forest fires and drought.

**Fig. 4.**
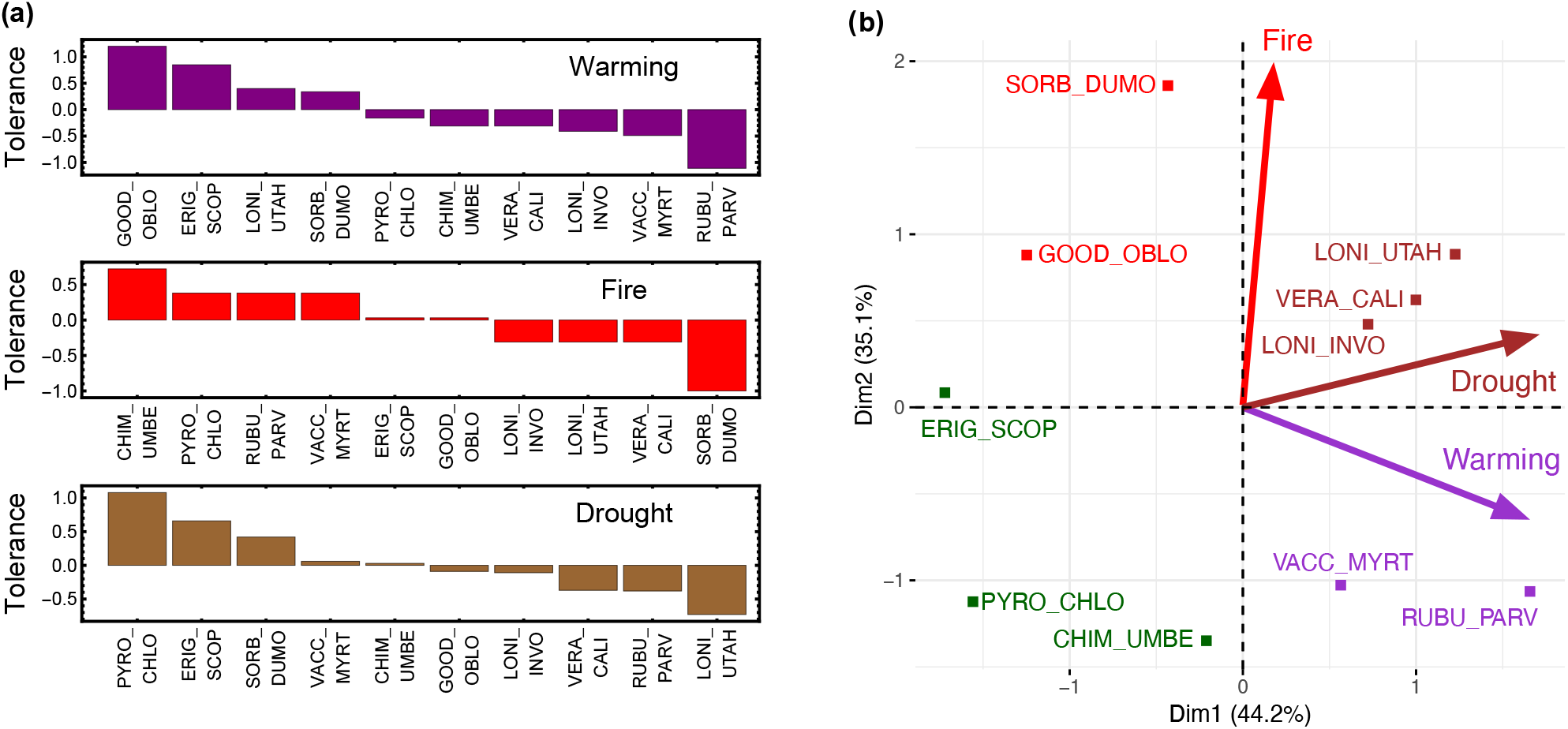
**(a)** Estimated species tolerances to the three environmental cues of climate change: warming, forest fire, and drought. See exact trait values in Table S6. **(b)** Estimated species intolerances (negative tolerances) to simultaneous and equal action of the warming, forest fire, and drought. For species codes see Table 1.

### Comparison to the historical data

Comparison of current species observations with historical data of 1974 and 1993 (23, 24) is shown in Table 2. Species sensitive to warming (according to our trait-based model: *Vaccinium myrtillus, Rubus parviflorus*, see Figure 4b) – have not become extinct. Even contrary, the abundance and the vigor of populations of *Rubus parviflorus* have increased – in comparison to observations in 1993. Note that in previous surveys, different metrics of abundance were used for some plants. Thus, *Vaccinium myrtillus* was previously reported as ‘occasional small populations’ (24), without counting locations and/or ramets. Conservatively interpreting the historical data, we assumed the changes in abundance of *Vaccinium* are not significant, however we point out the low vigor of population of this species. Fire-sensitive species (according to the model: *Sorbus dumosa* and *Goodyera oblongifolia)* decreased since 1993 both in abundance and vigor, as did most of drought-sensitive species, except *Veratrum californicum*, which was still common, robust, and dominating wet sites along springs to the exclusion of other species. *Chimaphila umbellata* was reported in 1993 as quite common, nevertheless we did not observe any individuals of these plants, only abundant populations of *Chimaphila maculata* along the Morse Canyon and Crest trails. We did not estimate the abundance of *Erigeron scopulinus* since plants reported in 1993 (at rock outcrops near Raspberry Peak; (24)) were outside the survey area.

**Table 2.** Number of species locations and number of individuals in 1974, 1993, 1994 (after Rattlesnake Fire), and in 2019. Population vigor (averaged per species) in 1993 and 2019. Changes in species abundance and vigor 1993–2019 are marked as red arrows (decreasing), green arrows (increasing), or white circles (no change). Abbreviations: occ. – occasionally, com. – common; part.surv. – partly survived.

| Species code | Number of locations/individuals |  |  |  | Changes<br>1993-2019 | Vigor |  | Changes<br>1993-2019 |
| --- | --- | --- | --- | --- | --- | --- | --- | --- |
|  | 1974 | 1993 | 1994 | 2019 |  | 1993 | 2019 |  |
| CHIM_UMBE | 1/20 | 10/220 | gone? | 0/0 | ▼ | 2 | - | - |
| ERIG_SCOP | 1/3 | 1/3 | gone | 0/0 | ? | 2 | - | - |
| GOOD_OBLO | 2/11 | 3/15 | part.surv. | 1/7 | ▼ | 2 | 1.5 | ▼ |
| LONI_INVO | 4/4 | 7/7 | no data | 2/6 | ▼ | - | 2.8 | - |
| LONI_UTAH | 24/24 | 21/21 | part.surv. | 7/18 | ▼ | 2 | 1.6 | ▼ |
| PYRO_CHLO | 1/3 | 1/4 | gone | 3/9 | ▲ | - | 1.8 | - |
| RUBU_PARV | occ. | 4/38 | gone? | 6/259 | ▲ | 2 | 2.5 | ▲ |
| SORB_DUMO | 1/1 | 4/4 | gone | 1/1 | ▼ | 2 | 1 | ▼ |
| VACC_MYRT | occ. | occ. | part.surv. | 5/90 | ○ | 1 | 1 | ○ |
| VERA_CALI | com. | com. | survived | 5/389 | ○ | 3 | 2.9 | ○ |

## Discussion

The comparison of the species abundance and vigor with historical data of 1974 and 1993 shows varied responses of the species to climate change in the Chiricahua Mountains. Such differences are also indicated by our plant-functional-trait-based model (see Figure 4b) showing that the chosen remnant boreal plant species have various life histories. Since 1993, the study area endured severe large forest fires and pressure from gradual climate change. We found that all plant species sensitive to fire, first, had increased in abundance (in 1975–1993) and then, evidently decreased in both abundance and vigor (in 1994–2019), as did also most of drought-sensitive species. It is explained by the local climate data: the period of 1975–1993 was characteristic with increased amount of precipitation (see Figure 2b) and with no large forest fires. Meantime, the pressure of the global warming remains controversial. One of the species most sensitive to warming – *Rubus parviflorus*, has increased in abundance and vigor. The second most warming-sensitive species – *Vaccinium myrtillus*, has remained more or less unchanged in terms of abundance and low vigor. We explain the low vigor of *Vaccinium* with the fact that this dwarf shrub needs deep snow cover during winters (33, 34), but the local snow depth decreased since 1994 (see Figure 2d) that could be both due to decrease in precipitation (*ρ* = 0.44, see Results) and due to increased exposure to sun and wind caused by two large forest fires. Lower vigor of *Vaccinium* may be occurring not because of any short climate trend, but possibly driven by a more general warming since the end of the Little Ice Age about 150 years ago (24). In fact, increasing temperatures tend to increase evaporation, which leads to more precipitation. However, the regional precipitation pattern could be different from the global trend (35) and needs careful consideration of all involved factors. The large forest fires are recognized as synergistic consequences of both the decrease in precipitation and the exclusion of fire as a forest management tool (8).

We did not find two of the three plant species, which appeared to be most resistant to all studied environmental cues according to our functional-trait-based model. Neither of these two species are appropriate for testing our hypothesis and were included in our study to be consistent with previous surveys. One of these species – *Chimaphila umbellata* subsp. *acuta* – in the study area, grows along the northern edge of its range (36) and, consequently, is not suitable for testing our hypothesis about the southern-edge-species persistence. Another missing plant – *Erigeron scopulinus* – is extremely rare (only two known locations in the Chiricahua Mountains (37)) and we did not encounter this plant because our survey was of limited extent (only 10 meters band along trails, following methodology) that was not appropriate for counting rare species. Also, this species does not necessarily need a cryic soil temperature (24). The good condition of *Veratrum californicum* we explain by the fact that this species is not on it’s southernmost geographic range limit in the Chiricahua Mountains as its distribution range extends into northern Mexico. In favor of the species persistence serves a proximity of permanent water sources as evidenced by findings of *Veratrum californicum*, as well as *Rubus parviflorus*, and the only individuals of *Sorbus dumosa* and *Lonicera involucrata*, which were encountered only in wet canyons. Such water sources could mitigate the adverse impacts of drought and forest fire and thus serve as a refugia (10). However, the wet canyons do not prevent invasion pressure from lower-elevation plant communities enhanced by climate warming.

While warming tolerance (sensitivity) is actively studied and discussed worldwide (38–41), such studies are practically limited by lack of consistent knowledge about plant functional traits. Particularly, data on tolerance to warming are missing in existing plant trait databases, and data on tolerances to fire and drought are available only for about 1.7% and 0.95% of all accepted taxonomic names of vascular plant species, respectively (27, 42). In this paper, we estimated the warming tolerance by using traits directly associated with temperature extremes for a given species in its natural range. Note that tolerance to warming is not strongly related to species maximum temperature tolerance, but rather is limited by indirect factors such as competition from thermophilic species. It is especially true for cryophilic species that rely on stress-tolerant strategy and are a little competitive. A more challenging but powerful approach would be to estimate warming tolerance from multiple soft traits such as plant parts characteristics or from ‘harder’ traits related to life-history strategies. For wider discussion of the warming tolerance assessment see (43). In all cases, the warming tolerance needs to be considered together with other environmental cues, when modeling warming impacts to a plant taxon. Of particular importance is to identify indicator species that are sensitive to temperature change but resistant to other environmental stress factors. Alternatively, to identify and study localities/regions, where climate warming is a dominant stress factor, for example, localities around a stable water supply. Note that species’ warming tolerances in this study are a practical measure of the species vulnerability (44), since we study remnant boreal plants with presumably high exposure to warming and that have limited adaptation capacity due to small populations isolated on a sky island in the arid matrix, which translates to low genetic diversity (45).

When analyzing historical and weather data, we encountered some challenges typical for the studied time period. During the last decades of the 20^th^ century, the technologies, scientific methods and metrics were drastically changed, leading to disparities in observations. These disparities are also widely propagated in climate models that extend and downscale the observations from weather stations. Thus, we had to conclude that reliable estimates of local temperature changes are unlikely since the weather observation methods were changing during the studied time period. New weather stations have made weather monitoring nearly continuous and potentially bias-free, but such data are available for the Chiricahua Mountains only since 2016. For further studies, it is reasonable to quantify local climate history by using core samples from local trees – as the most accessible, trustful and developed method (32). Using core samples, it should be noted that trees growing near a permanent water source could be less sensitive to drought, but trees in open spaces could be more exposed to temperature extremes. By analyzing core samples from trees exposed differently to environmental cues (stresses) of climate change, one could exclude combined actions of some of these stresses.

## Conclusions

We compared the historical and current abundance/vigor of ten remnant boreal plant species at high elevation in the Chiricahua Mountains referring to a 45-yrs time period (1974, 1993–1994 and 2019) and compared the field results with a functional-trait-based prediction. We observed that all species except two ones still occur in the study area, however five of them are in low vigor. We used functional traits of the surveyed plant species to model their tolerances to the three environmental stresses: warming, forest fire, and drought. Our model predicted that only *Rubus parviflorus* and *Vaccinium myrtillus* are selectively sensitive to warming, but other plant species are rather sensitive to fire, as well as to decrease in precipitation. Our comparison with the previous field surveys showed evident adverse impacts of large forest fires and decrease in precipitation in the study area since 1994. However, we did not find significant impact of climate warming on abundance and vigor of the surveyed species. Populations of the warming-sensitive species have not declined. *Rubus parviflorus* has increased in number and is in good vigor, but populations of *Vaccinium myrtillus* persist in the same abundance and vigor. In our work, we demonstrate how a combination of two basic ecological methods – field survey and modeling, can be used for better understanding and prediction of the ecological consequences of different environmental factors to support environmental management decisions, particularly the decisions related to climate change. Further combined studies are necessary to model climate-species interactions, to collect comprehensive spatio-temporal data on species abundance and vigor, and finally, to predict and prevent species and ecosystems losses due to climate change.

## ACKNOWLEDGEMENTS

We thank Isabel DeVito, Dylan Winkler and Frank Insana for field work, botanist Elaine Moisan for practical advice and plant identification, and SWRS administrator Alina Downer for kind support. We also appreciate the advice of Prof. Ursula Schuch and useful comments on this manuscript from Dr. Thomas Wohlgemuth.

## Supplementary Information

### Supplementary Note 1

**Table S1.** List of trails surveyed.

| No | Trail name and number | Length, km |
| --- | --- | --- |
| 1 | Crest Trail #270 | 9.709 |
| 2 | Greenhouse Trail #248 (upper part until Winn Falls) | 3.082 |
| 3 | Anita Park Trail #359 | 0.323 |
| 4 | Booger Spring Trail #347 | 0.276 |
| 5 | Tub Spring Trail (0.5 km along Centella Trail #334) | 0.315 |
| 6a | Monte Vista Lookout Trail, consisting from:<br>a) Morse Canyon Trail #43 | 3.404 |
| 6b | b) Crest Trail - Junction Saddle to Monte Vista Peak #270B | 3.411 |
| Total: |  | 20.520 |

### Supplementary Note 2

**Table S2.** Vigor of plant species accessed according the three vigor classes.

| Class | Species vigor | Description |
| --- | --- | --- |
| 1 | Low vigor | No fruits, no flowers, weak robustness, may have symptoms of diseases/herbivory/insects/parasitism |
| 2 | Medium vigor | May have some flowers/fruits but have symptoms of diseases/herbivory/insects/parasitism; or – healthy but not flowering/ fruiting |
| 3 | High vigor | Healthy, strong, flowering/fruiting, no symptoms of diseases/herbivory/ insects/parasitism |

### Supplementary Note 3

**Table S3.** Species’ habitat fire severity assessed according (Keeley, 2009).

| Class | Fire severity | Description |
| --- | --- | --- |
| 1 | Unburned | Plants green and unaltered, no direct effect from heat |
| 2 | Scorched | Unburned but plants exhibit leaf loss from radiated heat |
| 3 | Light | Canopy trees with green needles although stems scorched Surface litter, mosses and herbs charred or consumed Soil organic layer largely intact and charring limited to a few mm depth |
| 4 | Moderate or severe surface burn | Trees with some canopy cover killed, but needles not consumed All understory plants charred or consumed Fine dead twigs on soil surface consumed and logs charred Pre-fire soil organic layer largely consumed |
| 5 | Deep burning or crown fire | Canopy trees killed and needles consumed Surface litter of all sizes and soil organic layer largely consumed White ash deposition and charred organic matter to several cm depth |

## Supplementary Note 4: Detailed observations of the ten boreal species studied

All photos are taken by authors during the survey, in September-October 2019.

1. ***Chimaphila umbellata*** was not observed in 2019. However, there were about hundred individuals of *Chimaphila maculata* along the Morse Canyon and Crest trails (see Figure S1). *C. umbellata* was reported in 1993 as quite common (more than 200 individuals), but this observation was supported only by one record in herbarium collections (SEINet Portal Network, 2020). We suppose that this species, perhaps, might have been confused in the previous surveys with *C. maculata*.

**Fig. S1.**
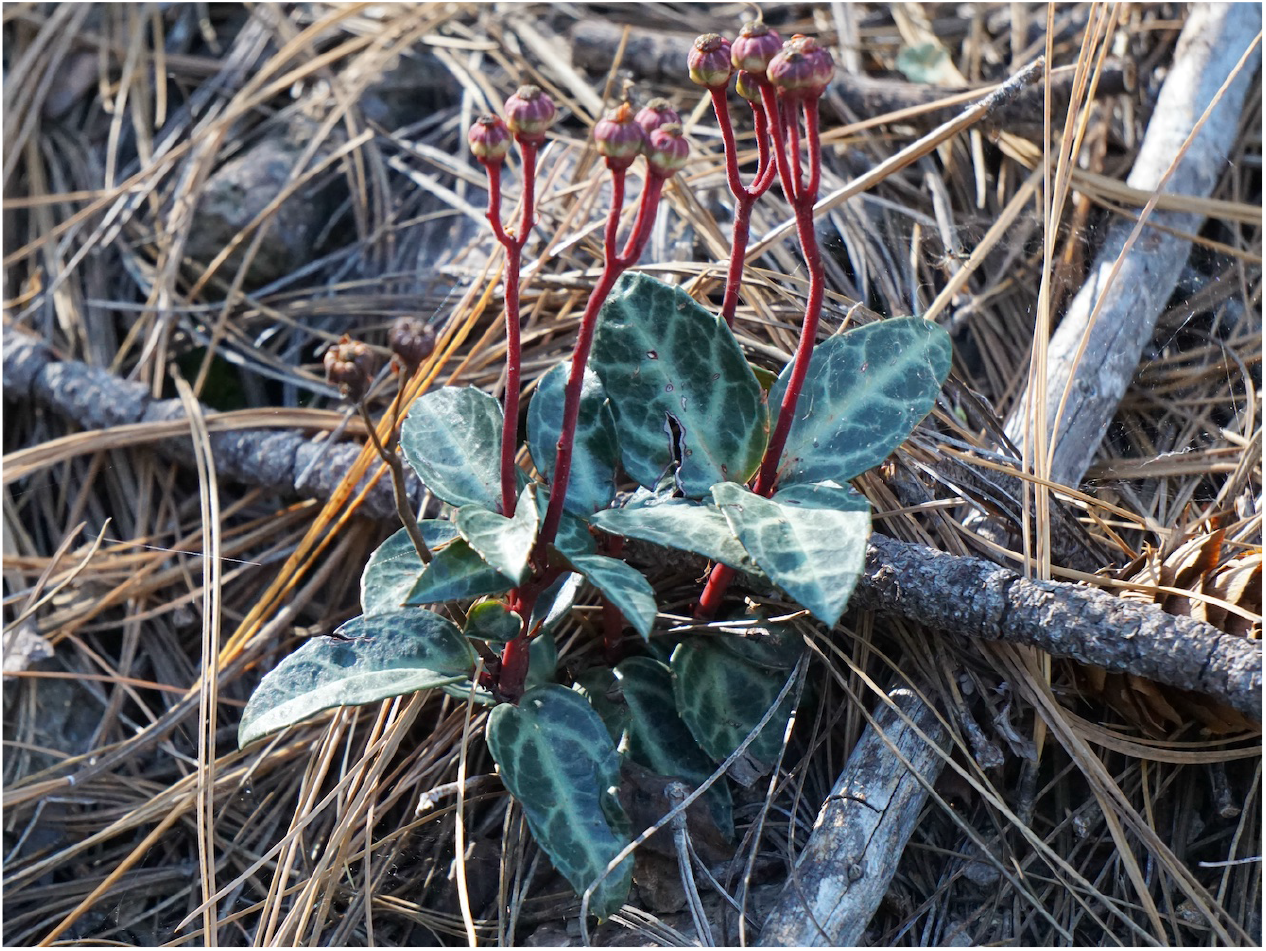
*Chimaphila maculata* on the Morse Canyon trail, Chiricahua Mountains, Arizona, October 2019.

2. ***Erigeron scopulinus*** was not observed in 2019, since we surveyed only 10 meters band of trails and did not surveyed rocks that were more distant from trails. Only two natural locations of this species in the Chiricahua Mountains are known from other studies (Nesom and Roth 1981). Meantime, the species is spread by commercial plant nurseries as a decorative alpine plant.

---

3. ***Goodyera oblongifolia*** was found in one plot along Crest Trail near Booger Spring, at the elevation of 2841 meters a.s.l., on dry soil. The population had 7 physically isolated small patches, which we counted as 7 separate ramets (altogether 38 shoots). Four of the shoots were flowering. However, the shoots were small (leaves only 2–4 cm long) and the population presumably suffered from drought and trampling since the plants grew very close to trail. This location of *Goodyera* was mentioned also in 1993 and 1994. We did not find other two historical locations of this orchid. Fire severity in the plot was estimated as light. Dominant tree species: Engelmann spruce *(Picea engelmannii)*, Douglas fir *(Pseudotsuga menziesii)*. Herbarium specimen collected for the SWRS Herbarium.

**Fig. S2.**
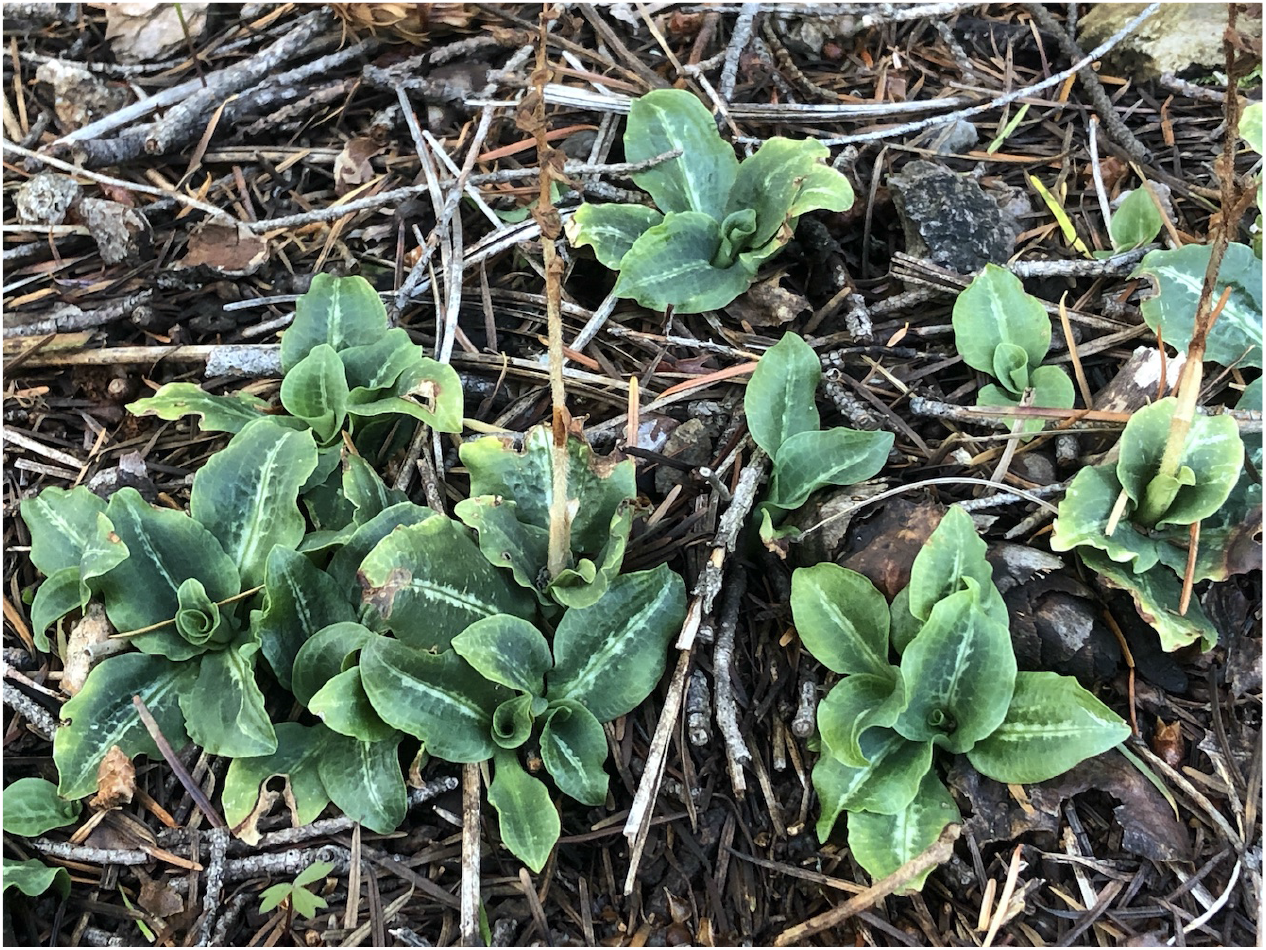
*Goodyera oblongifolia* on the Crest Trail near Booger Spring, Chiricahua Mountains, Arizona, September 2019.

4. ***Lonicera involucrata*** was found in two separate plots (two locations) along Cima Creek (Greenhouse Trail) above Winn Falls, at the elevations of 2663 and 2646 meters a.s.l., respectively. Both individuals were likely two of the seven ones, observed in 1993. Both individuals were strong (2-3 meters high), healthy and abundantly flowering/fruiting. In one location, also 4 new healthy individuals up to 1-meter high were found about 2-3 meters away from the old shrub. Fire severity was estimated as light in both locations. Dominant tree species: white fir *(Abies concolor*), quaking aspen *(Populus tremuloides*), Rocky Mountain maple *(Acer glabrum)*, Douglas fir, Engelmann spruce, Arizona pine *(Pinus arizonica)*, Gambel oak *(Quercus gambelii*). Herbarium specimen collected for the SWRS Herbarium.

**Fig. S3.**
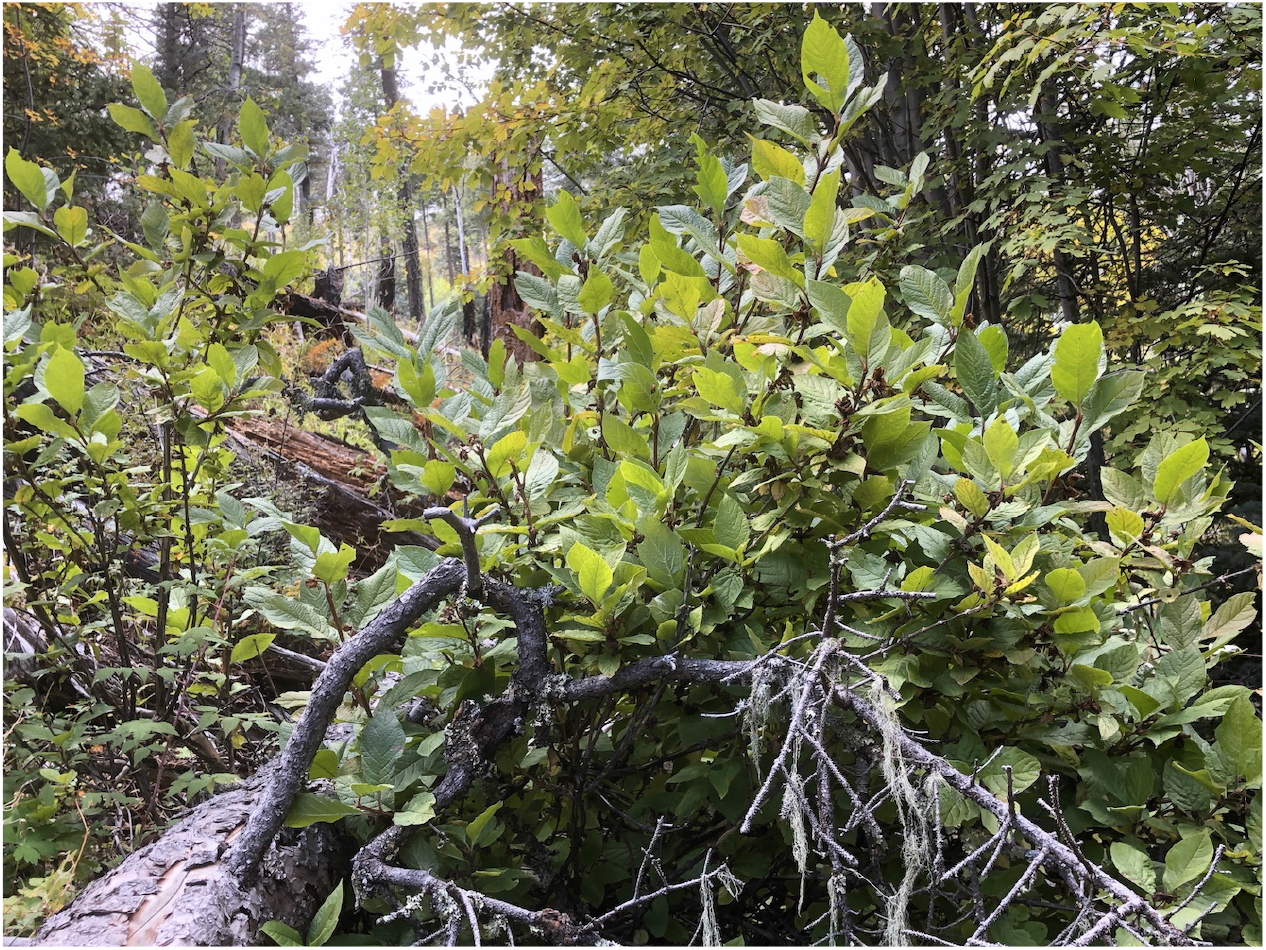
*Lonicera involucrata* on the Greenhouse Trail along Cima Creek, Chiricahua Mountains, Arizona, October 2019.

5. ***Lonicera involucrata*** on the Greenhouse Trail along Cima Creek, Chiricahua Mountains, Arizona, October 2019.*Lonicera utahensis* was found at 9 plots (clustered in 7 locations) scattered along all trails and at wide range of elevations: from lowest (2607 meters) to highest (2910 meters). However, most of individuals (at 6 plots) were weak, not reaching 0.5-meter height and not flowering/fruiting. Fire severity varied from light to deep. Dominant tree species: white fir, aspen, Rocky Mountain maple, Douglas fir, Engelmann spruce, Arizona pine, Gambel oak.

**Fig. S4.**
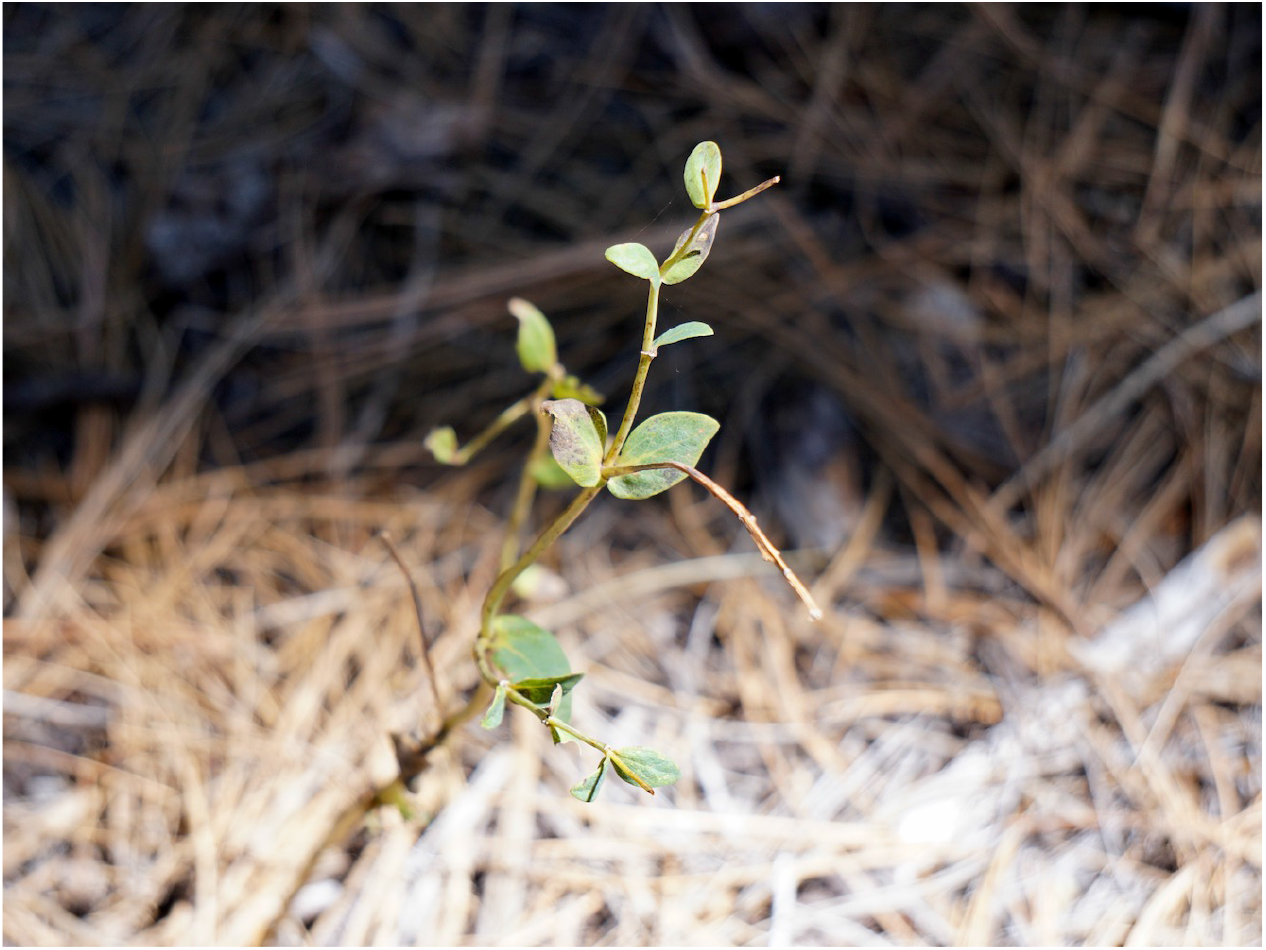
*Lonicera utahensis* on the Monte Vista Lookout Trail, Chiricahua Mountains, Arizona, October 2019.

6. ***Pyrola chlorantha*** grew at three plots along Morse Canyon Trail at the elevation of 2191-2427 meters a.s.l., on the north or northwest slope. Number of separated ramets varied from 1 to 4 per plot (with total shoot number 21). Vigor of species was estimated as medium. About half of shoots were flowering. *Pyrola chlorantha* shared the trail with much more abundant *Chimaphila maculata*. Fire severity estimated as light in all three plots. Dominant tree species: Arizona pine. Herbarium specimen collected for the SWRS Herbarium.

**Fig. S5.**
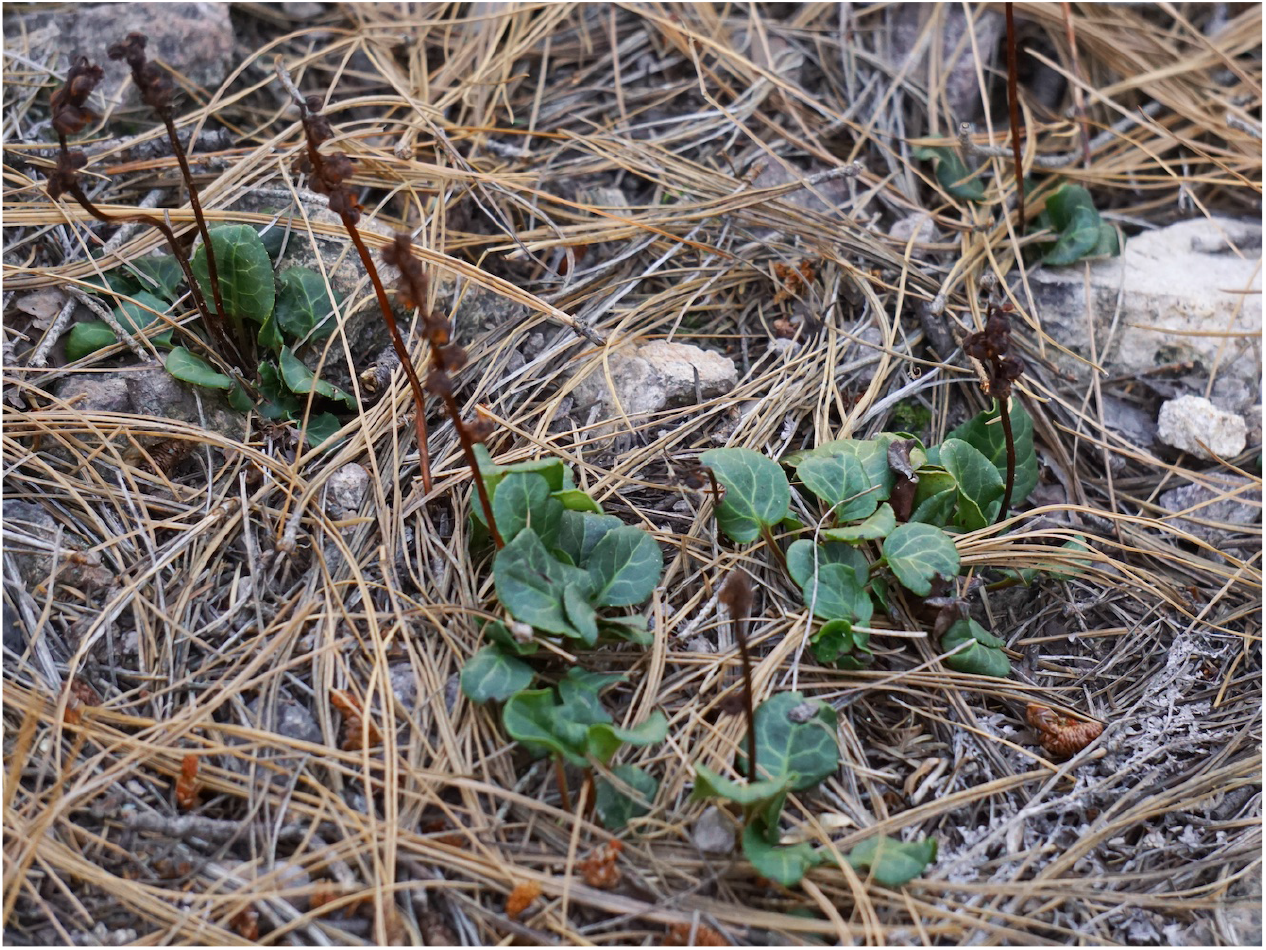
*Pyrola chlorantha* on the Morse Canyon trail, Chiricahua Mountains, Arizona, October 2019.

7. ***Rubus parviflorus*** found frequent (22 plots) along the Greenhouse Trail, in the vicinity of Cima Creek, while along the Crest Trail a similar *R. neomexicanus* was quite common. Shoots were mostly robust; in 11 plots individuals were flowering/fruiting. Vigor of population estimated as medium to high. Elevation fluctuated from 2622 to 2706 meters a.s.l. Found mostly on a bottom of the canyon on both banks of the creek, also along the trail (on the south or southwest slope). Fire severity varied from scorched to moderate. Dominant tree species: white fir, aspen, Rocky Mountain maple, Douglas fir, Engelmann spruce, Arizona pine, Gambel oak. Herbarium specimen collected for the SWRS Herbarium.

**Fig. S6.**
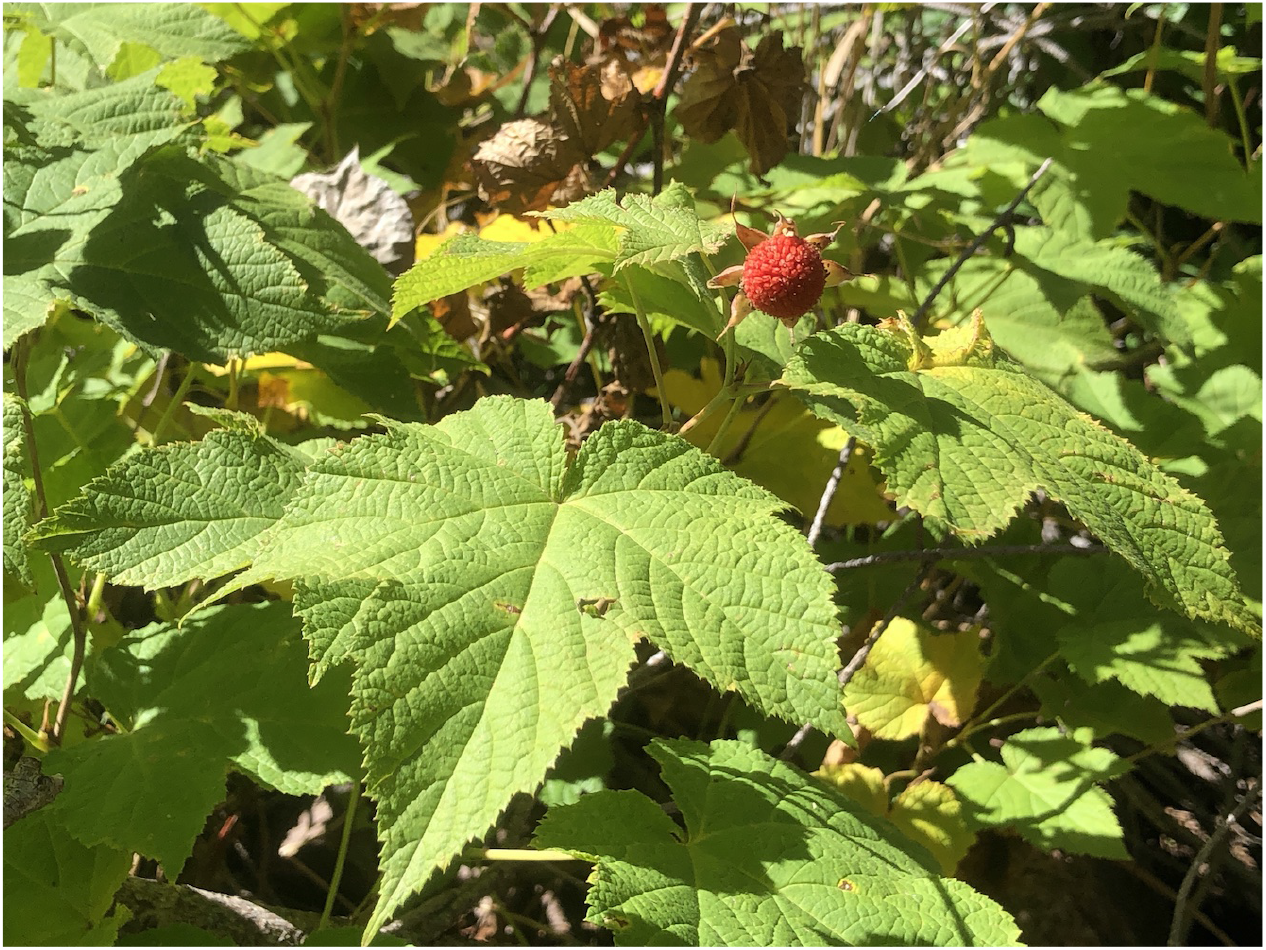
*Rubus parviflorus* on the Greenhouse Trail along Cima Creek, Chiricahua Mountains, Arizona, October 2019.

8. ***Sorbus dumosa*** found only one individual at one location on moist soil on the bank of Cima Creek. The shrub was about 2 meters high, no flowering, no fruiting. Elevation 2659 meters a.s.l. Fire severity estimated as light. Dominant tree species: aspen, Douglas fir, Arizona pine, Gambel oak. We did not find individuals neither above Cima Cabine, nor near Raspberry Peak, which were most likely gone after large forest fires. Herbarium specimen collected for the SWRS Herbarium.

**Fig. S7.**
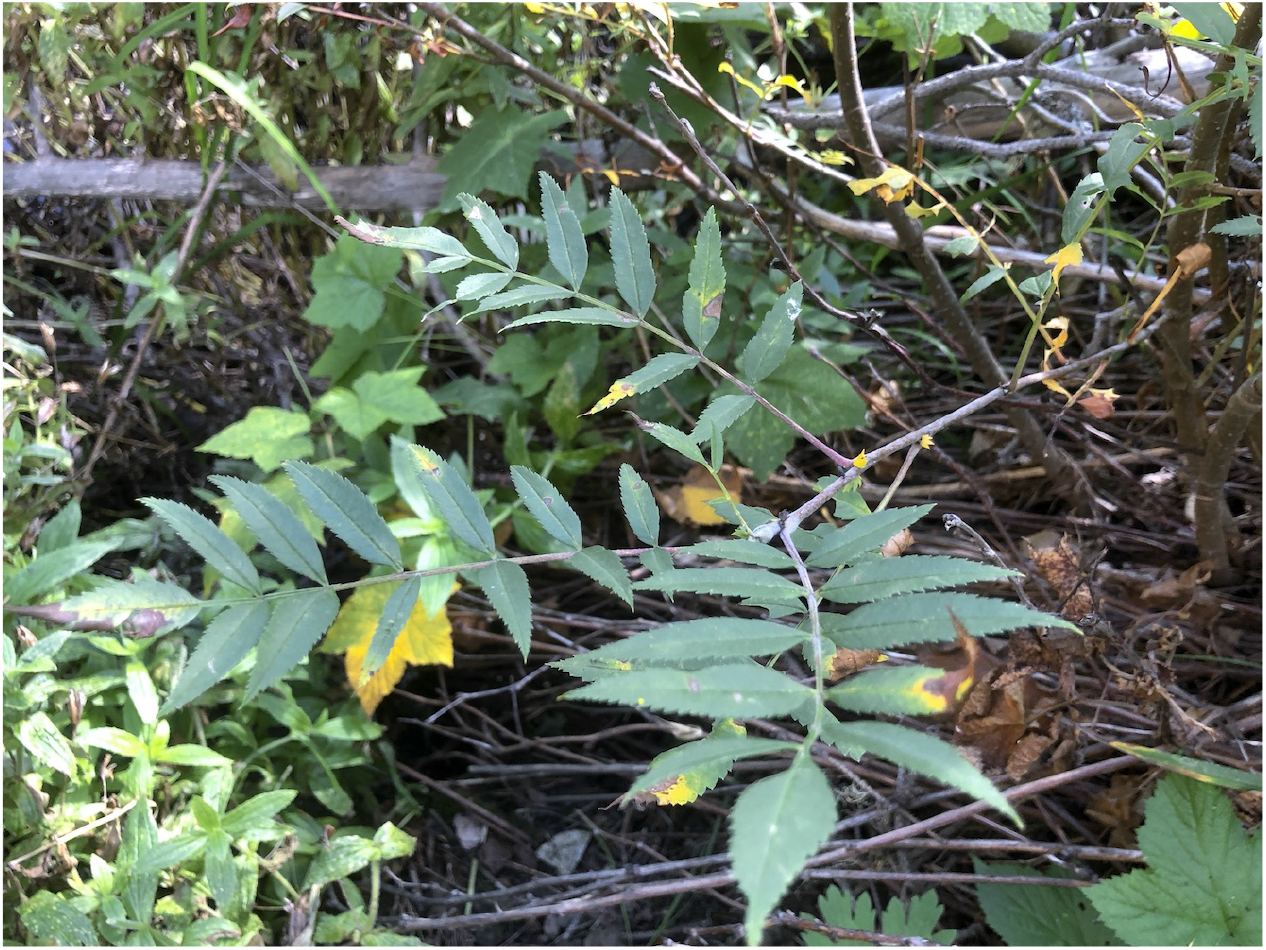
*Sorbus dumosa* on the Greenhouse Trail along Cima Creek, Chiricahua Mountains, Arizona, October 2019.

9. *Vaccinium myrtillus* Found at 11 plots (clustered in 5 locations) in open woods and only at the upmost elevations (above 2800 meters a.s.l.). Despite the relative abundance, the shoots were weak and sparse and some of them had only a few leaves. No flowering/fruiting. Fire severity estimated as moderate to severe, in some locations – deep. Dominant tree species: aspen, Douglas fir, Arizona pine. Herbarium specimen collected for the SWRS Herbarium.

**Fig. S8.**
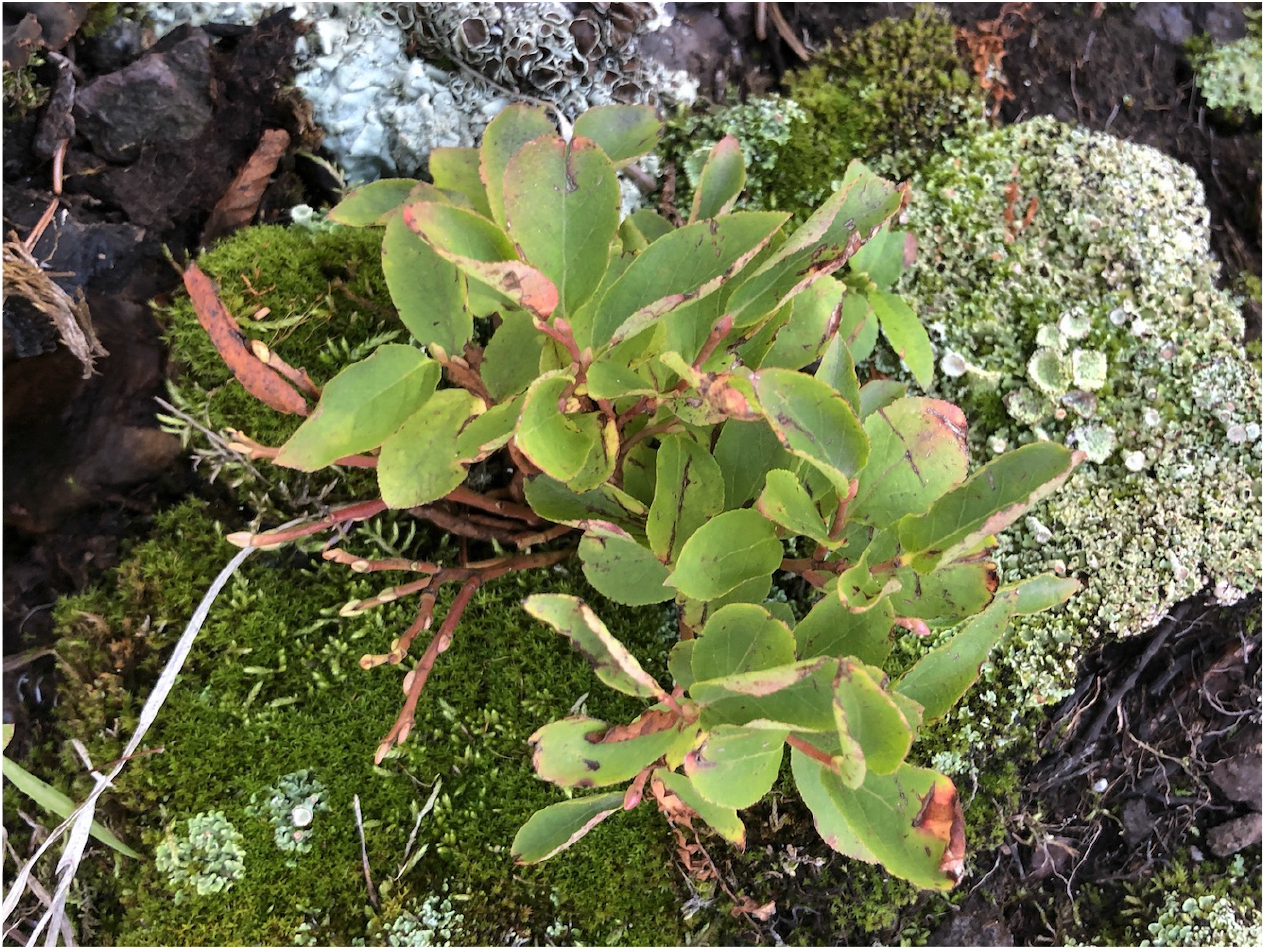
Vaccinium myrtillus on the Crest Trail, Chiricahua Mountains, Arizona, September 2019.

10. ***Veratrum californicum*** Common along Anita Spring, Booger Spring, Tub Spring, and Cima Creek (starting from Cima Cabin to the east). Shoots were robust, most of them flowering/fruiting. Vigor of population estimated as high. Fire severity estimated as scorched to light. Dominant tree species: aspen, white fir, Rocky Mountain maple, Douglas fir, Engelmann spruce, Arizona pine. This species fits well inside its geographical range and seems feeling quite well in the Chiricahua Mountains. Herbarium specimen collected for the SWRS Herbarium.

**Fig. S9.**
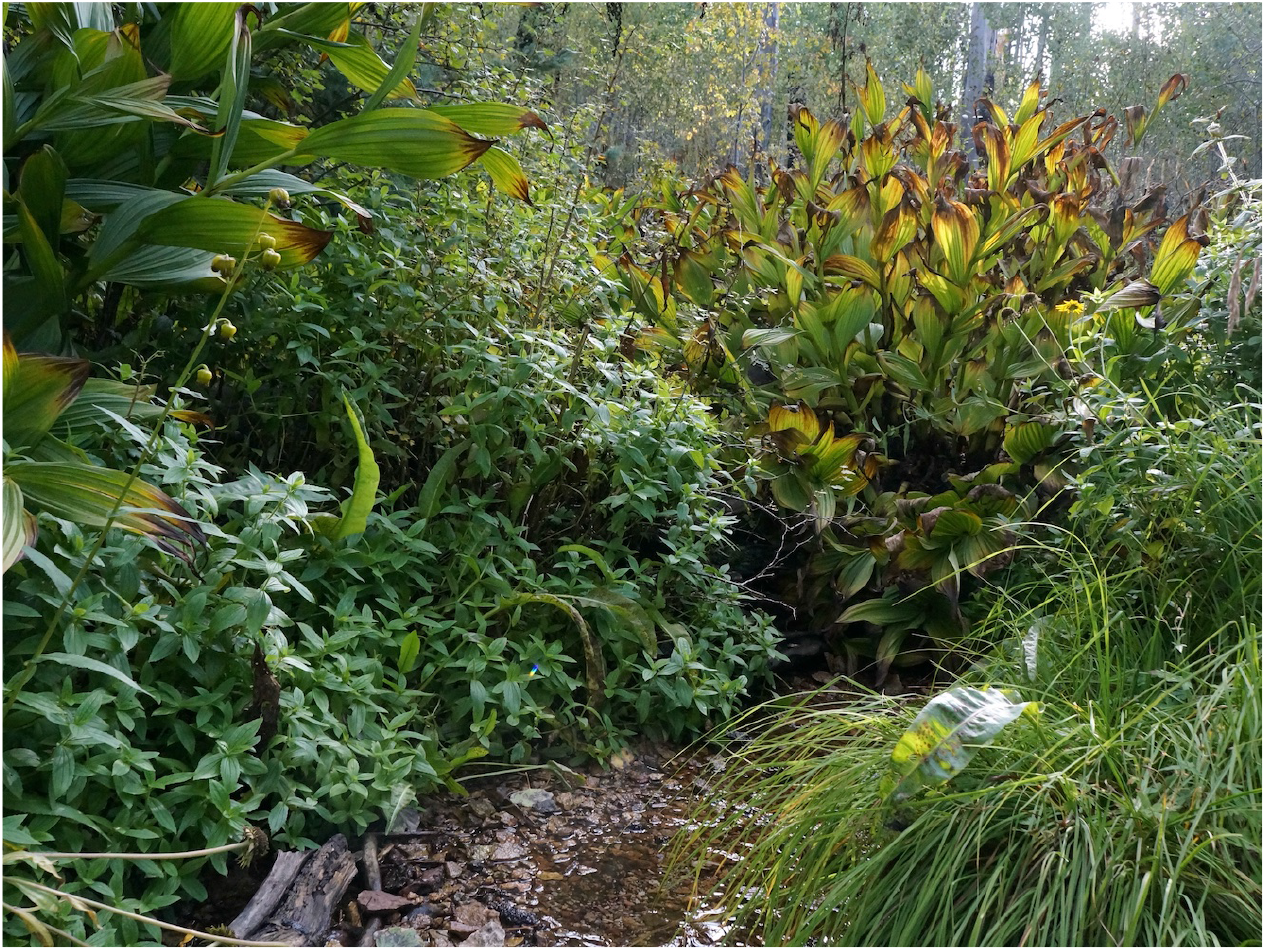
*Veratrum californicum* near Tub Spring, Chiricahua Mountains, Arizona, September 2019.

## Supplementary Note 5

**Table S4.**
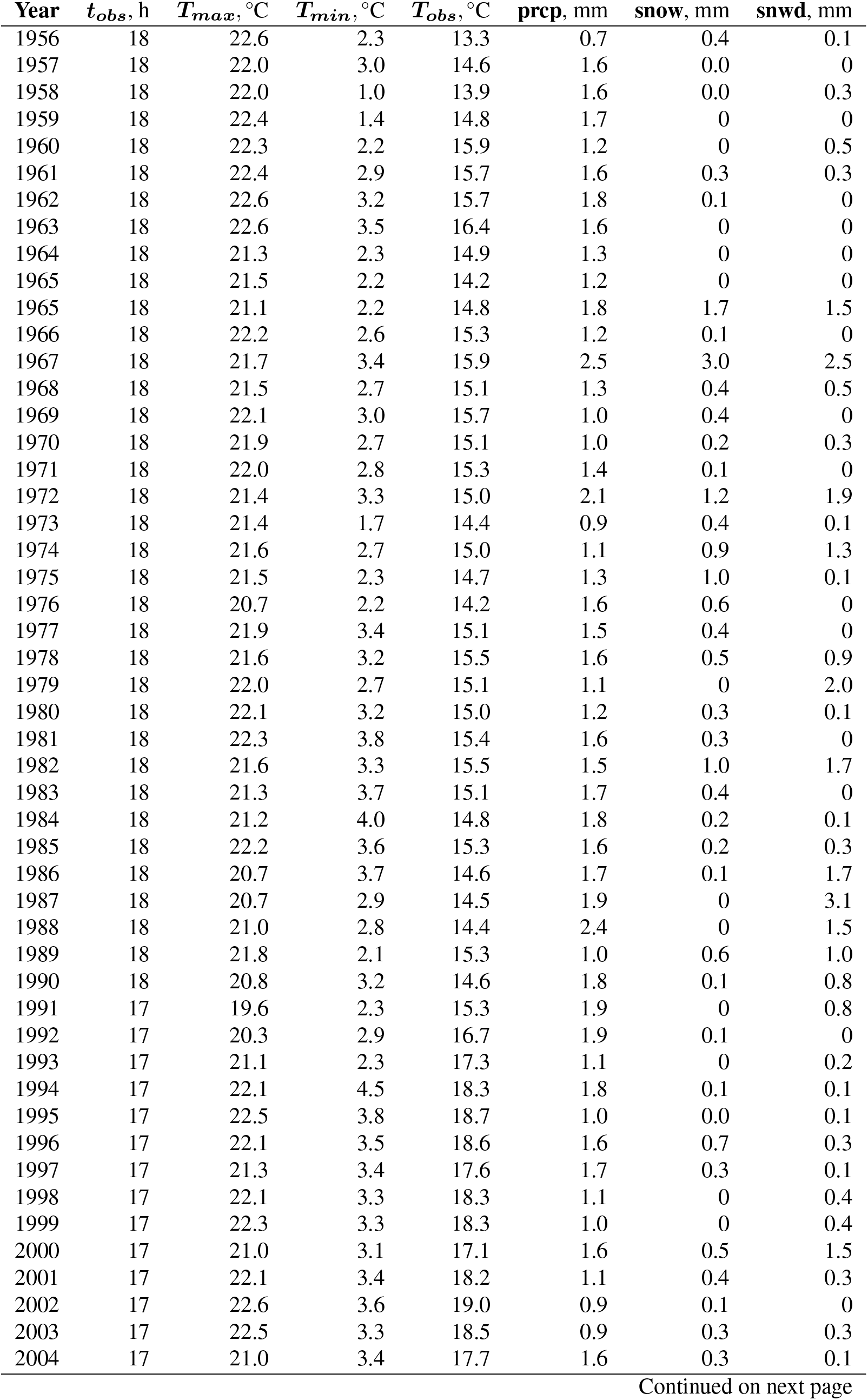
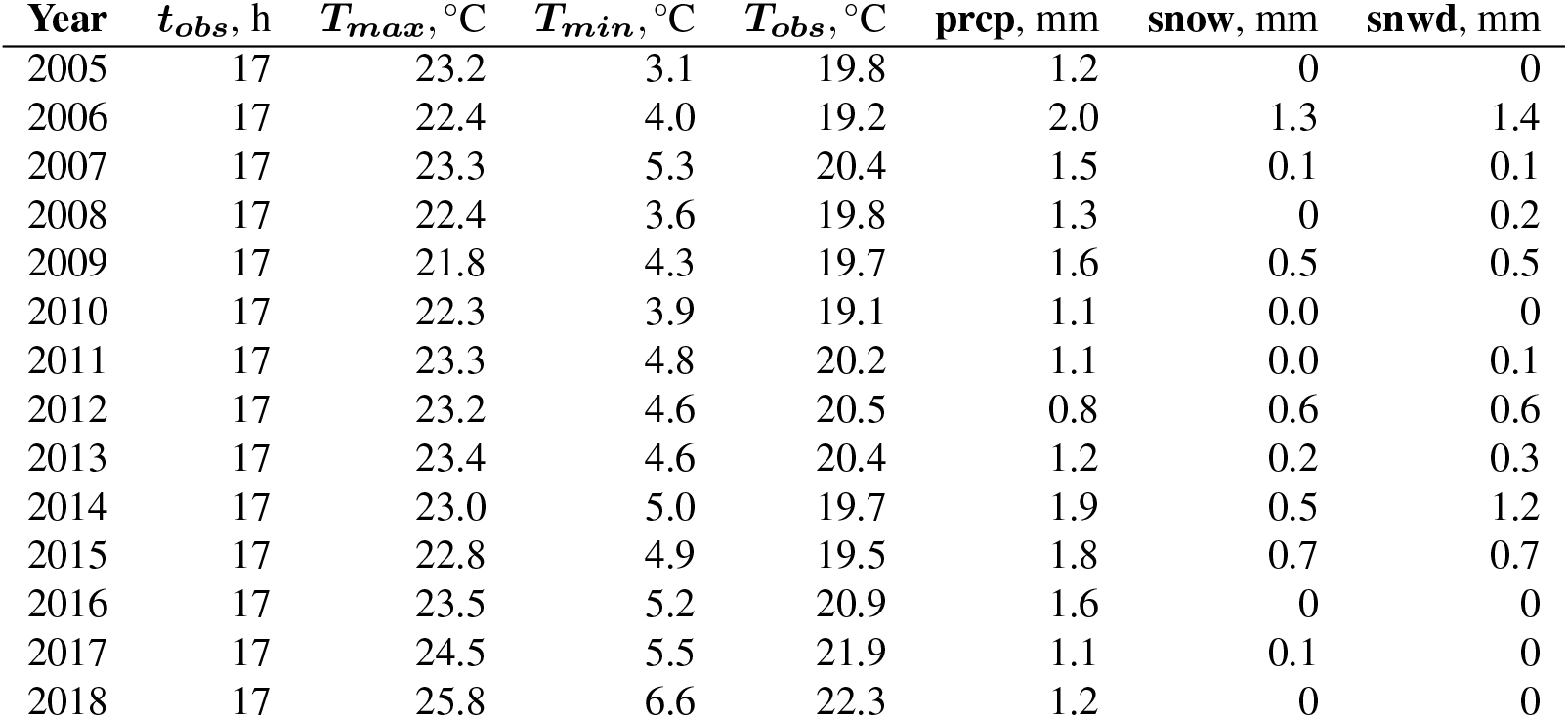
Local weather data of the PORTAL 4 SW weather station, daily observations averaged per year. *T_max_*: maximum temperature; *T_min_*: minimum temperature; *T_obs_*: temperature at the time of observation; *t*_obs_: time of observation; **prcp**: precipitation; **snow**: snowfall; **snwd**: snow depth.

| Year | $t_{obs}$, h | $T_{max}$, °C | $T_{min}$, °C | $T_{obs}$, °C | prcp, mm | snow, mm | snwd, mm |
| --- | --- | --- | --- | --- | --- | --- | --- |
| 1956 | 18 | 22.6 | 2.3 | 13.3 | 0.7 | 0.4 | 0.1 |
| 1957 | 18 | 22.0 | 3.0 | 14.6 | 1.6 | 0.0 | 0 |
| 1958 | 18 | 22.0 | 1.0 | 13.9 | 1.6 | 0.0 | 0.3 |
| 1959 | 18 | 22.4 | 1.4 | 14.8 | 1.7 | 0 | 0 |
| 1960 | 18 | 22.3 | 2.2 | 15.9 | 1.2 | 0 | 0.5 |
| 1961 | 18 | 22.4 | 2.9 | 15.7 | 1.6 | 0.3 | 0.3 |
| 1962 | 18 | 22.6 | 3.2 | 15.7 | 1.8 | 0.1 | 0 |
| 1963 | 18 | 22.6 | 3.5 | 16.4 | 1.6 | 0 | 0 |
| 1964 | 18 | 21.3 | 2.3 | 14.9 | 1.3 | 0 | 0 |
| 1965 | 18 | 21.5 | 2.2 | 14.2 | 1.2 | 0 | 0 |
| 1965 | 18 | 21.1 | 2.2 | 14.8 | 1.8 | 1.7 | 1.5 |
| 1966 | 18 | 22.2 | 2.6 | 15.3 | 1.2 | 0.1 | 0 |
| 1967 | 18 | 21.7 | 3.4 | 15.9 | 2.5 | 3.0 | 2.5 |
| 1968 | 18 | 21.5 | 2.7 | 15.1 | 1.3 | 0.4 | 0.5 |
| 1969 | 18 | 22.1 | 3.0 | 15.7 | 1.0 | 0.4 | 0 |
| 1970 | 18 | 21.9 | 2.7 | 15.1 | 1.0 | 0.2 | 0.3 |
| 1971 | 18 | 22.0 | 2.8 | 15.3 | 1.4 | 0.1 | 0 |
| 1972 | 18 | 21.4 | 3.3 | 15.0 | 2.1 | 1.2 | 1.9 |
| 1973 | 18 | 21.4 | 1.7 | 14.4 | 0.9 | 0.4 | 0.1 |
| 1974 | 18 | 21.6 | 2.7 | 15.0 | 1.1 | 0.9 | 1.3 |
| 1975 | 18 | 21.5 | 2.3 | 14.7 | 1.3 | 1.0 | 0.1 |
| 1976 | 18 | 20.7 | 2.2 | 14.2 | 1.6 | 0.6 | 0 |
| 1977 | 18 | 21.9 | 3.4 | 15.1 | 1.5 | 0.4 | 0 |
| 1978 | 18 | 21.6 | 3.2 | 15.5 | 1.6 | 0.5 | 0.9 |
| 1979 | 18 | 22.0 | 2.7 | 15.1 | 1.1 | 0 | 2.0 |
| 1980 | 18 | 22.1 | 3.2 | 15.0 | 1.2 | 0.3 | 0.1 |
| 1981 | 18 | 22.3 | 3.8 | 15.4 | 1.6 | 0.3 | 0 |
| 1982 | 18 | 21.6 | 3.3 | 15.5 | 1.5 | 1.0 | 1.7 |
| 1983 | 18 | 21.3 | 3.7 | 15.1 | 1.7 | 0.4 | 0 |
| 1984 | 18 | 21.2 | 4.0 | 14.8 | 1.8 | 0.2 | 0.1 |
| 1985 | 18 | 22.2 | 3.6 | 15.3 | 1.6 | 0.2 | 0.3 |
| 1986 | 18 | 20.7 | 3.7 | 14.6 | 1.7 | 0.1 | 1.7 |
| 1987 | 18 | 20.7 | 2.9 | 14.5 | 1.9 | 0 | 3.1 |
| 1988 | 18 | 21.0 | 2.8 | 14.4 | 2.4 | 0 | 1.5 |
| 1989 | 18 | 21.8 | 2.1 | 15.3 | 1.0 | 0.6 | 1.0 |
| 1990 | 18 | 20.8 | 3.2 | 14.6 | 1.8 | 0.1 | 0.8 |
| 1991 | 17 | 19.6 | 2.3 | 15.3 | 1.9 | 0 | 0.8 |
| 1992 | 17 | 20.3 | 2.9 | 16.7 | 1.9 | 0.1 | 0 |
| 1993 | 17 | 21.1 | 2.3 | 17.3 | 1.1 | 0 | 0.2 |
| 1994 | 17 | 22.1 | 4.5 | 18.3 | 1.8 | 0.1 | 0.1 |
| 1995 | 17 | 22.5 | 3.8 | 18.7 | 1.0 | 0.0 | 0.1 |
| 1996 | 17 | 22.1 | 3.5 | 18.6 | 1.6 | 0.7 | 0.3 |
| 1997 | 17 | 21.3 | 3.4 | 17.6 | 1.7 | 0.3 | 0.1 |
| 1998 | 17 | 22.1 | 3.3 | 18.3 | 1.1 | 0 | 0.4 |
| 1999 | 17 | 22.3 | 3.3 | 18.3 | 1.0 | 0 | 0.4 |
| 2000 | 17 | 21.0 | 3.1 | 17.1 | 1.6 | 0.5 | 1.5 |
| 2001 | 17 | 22.1 | 3.4 | 18.2 | 1.1 | 0.4 | 0.3 |
| 2002 | 17 | 22.6 | 3.6 | 19.0 | 0.9 | 0.1 | 0 |
| 2003 | 17 | 22.5 | 3.3 | 18.5 | 0.9 | 0.3 | 0.3 |
| 2004 | 17 | 21.0 | 3.4 | 17.7 | 1.6 | 0.3 | 0.1 |

**Table S4 – continued from previous page**
| **Year** | **$t_{obs}$, h** | **$T_{max}$, °C** | **$T_{min}$, °C** | **$T_{obs}$, °C** | **prcp, mm** | **snow, mm** | **snwd, mm** |
| --- | --- | --- | --- | --- | --- | --- | --- |
| 2005 | 17 | 23.2 | 3.1 | 19.8 | 1.2 | 0 | 0 |
| 2006 | 17 | 22.4 | 4.0 | 19.2 | 2.0 | 1.3 | 1.4 |
| 2007 | 17 | 23.3 | 5.3 | 20.4 | 1.5 | 0.1 | 0.1 |
| 2008 | 17 | 22.4 | 3.6 | 19.8 | 1.3 | 0 | 0.2 |
| 2009 | 17 | 21.8 | 4.3 | 19.7 | 1.6 | 0.5 | 0.5 |
| 2010 | 17 | 22.3 | 3.9 | 19.1 | 1.1 | 0.0 | 0 |
| 2011 | 17 | 23.3 | 4.8 | 20.2 | 1.1 | 0.0 | 0.1 |
| 2012 | 17 | 23.2 | 4.6 | 20.5 | 0.8 | 0.6 | 0.6 |
| 2013 | 17 | 23.4 | 4.6 | 20.4 | 1.2 | 0.2 | 0.3 |
| 2014 | 17 | 23.0 | 5.0 | 19.7 | 1.9 | 0.5 | 1.2 |
| 2015 | 17 | 22.8 | 4.9 | 19.5 | 1.8 | 0.7 | 0.7 |
| 2016 | 17 | 23.5 | 5.2 | 20.9 | 1.6 | 0 | 0 |
| 2017 | 17 | 24.5 | 5.5 | 21.9 | 1.1 | 0.1 | 0 |
| 2018 | 17 | 25.8 | 6.6 | 22.3 | 1.2 | 0 | 0 |

## Supplementary Note 6

**Table S5.** Exact values of species tolerances to warming, fire, and drought.

| **Species code** | **Tolerance to** |  |  | **Functional Group** |
| --- | --- | --- | --- | --- |
|  | **warming** | **fire** | **drought** |  |
| CHIM_UMBE | 0.12 | 0.72 | 0.03 | 4 |
| ERIG_SCOP | 0.77 | 0.03 | 0.66 | 4 |
| GOOD_OBLO | 1.29 | 0.03 | -0.09 | 3 |
| LONI_INVO | -0.39 | -0.31 | -0.11 | 2 |
| LONI_UTAH | -0.06 | -0.31 | -0.73 | 2 |
| PYRO_CHLO | 0.05 | 0.38 | 1.08 | 4 |
| RUBU_PARV | -0.98 | 0.38 | -0.38 | 1 |
| SORB_DUMO | 0.07 | -1.00 | 0.42 | 3 |
| VACC_MYRT | -0.56 | 0.38 | 0.06 | 1 |
| VERA_CALI | -0.31 | -0.31 | -0.37 | 2 |

